# Investigating VCAM-1 Targeted Nanoparticles and Annexin A1 Therapy using Dysfunctional-endothelium-on-a-chip

**DOI:** 10.1101/2021.05.09.443301

**Authors:** Salime Bazban-Shotorbani, Felicity Gavins, Martin Dufva, Nazila Kamaly

## Abstract

Atherosclerosis is an inflammation-driven disease of the arteries and one of the leading causes of global mortality. The initial pathological stage in atherosclerosis is dysfunctional endothelium (Dys-En), which results in loss of adherens-junctions between cells, thus enhancing permeability. Not only the enhanced permeability of Dys-En can be used as a nanoparticle targeting mechanism, but also the normalization and restoration of this phenomenon can be utilized as a potent anti-atherosclerotic therapy. This study aimed to recruit a robust biomicrofluidic model of Dys-En for 1) nanoparticle screening and 2) normalization assessments. The developed Dys-En-on-a-chip could successfully mimic the atherosclerotic flow condition, enhanced permeability, formation of actin stress fibers, and overexpression of vascular cell adhesion molecule 1 (VCAM-1), which are known as hallmarks of a Dys-En. The screening of VCAM-1 targeting nanoparticles with variable biophysicochemical properties showed that nanoparticle size plays the main role in nanoparticle targeting, and the design of nanoparticles in the range of 30-60 nm can highly increase their targeting to Dys-En. Moreover, treatment of Dys-En-on-a-chip with Annexin A1, as a novel pro-resolving mediator, resulted in restoration of adherens-junctions and normalization of the barrier integrity. This data validates the use of biomicrofluidic models for investigating treatment regimens with biologics and to identify optimal nanoparticle properties for effective atherosclerotic plaque targeting.

## 1. Introduction

Cardiovascular diseases (CVD) are the leading cause of death worldwide, causing up to 18 million deaths annually ^1^. CVD is estimated to remain the main cause of mortality, resulting in 24 million global deaths per year by 2030 ^2^. The dominant cause of CVD is atherosclerosis, which is an inflammation-driven disease of the arteries ^3^. This number-one killer disease results in the formation of atherosclerotic plaques on arterial walls and can eventually cause thrombosis, myocardial infarction, and ischemic stroke ^4,5^. Despite the growing need for new and improved atherosclerosis therapies, the FDA approvals for new atherosclerotic therapeutics decreased during the last decade ^2^. One of the reasons behind this problem is the lack of predictive and effective disease models to test novel therapeutics prior to clinical studies ^6,7^. In vivo models are lengthy, costly, labor-intensive, ethically problematic, and poor predictors of human responses ^7^. On the other hand, conventional in vitro models cannot fully replicate the pathophysiological condition of this disease ^8^. A suitable vascular disease model should be able to mimic the physiological condition of a vessel (e.g. vessel dimensions, flow rate, shear stress, etc.) and the relevant pathological condition of the disease. Therefore, to overcome the shortcomings of the conventional in vitro and in vivo models, biomicrofluidic vascular models have recently gained a significant attention ^8–11^.

In order to develop a robust biomimetic and biomicrofluidic model for atherosclerotic research, we focused on dysfunctional endothelium (Dys-En) as one of the main characteristics of atherosclerosis. In a healthy vessel, nutrients and oxygen are supplied to the intima by their diffusion from the lumen. Furthermore, a microvasculature network called *vasa vasorum*, supplies oxygen and nutrients to the outer compartment of the vessel walls ^12,13^ (**Figure 1A**). Due to the plaque formation in an atherosclerotic vessel, the intima thickens, resulting in an insufficient and impair diffusion of oxygen and nutrients from the lumen to the intima. To restore oxygen and nutrients supply, neovascularization occurs and the *vasa vasorum* sprout into the atherosclerotic plaque ^14,15^. In contrast to a healthy endothelium (En), the endothelium on these intraplaque neovessels is inflamed and Dys-En ^16^ (**Figure 1B**).

**Figure 1.**
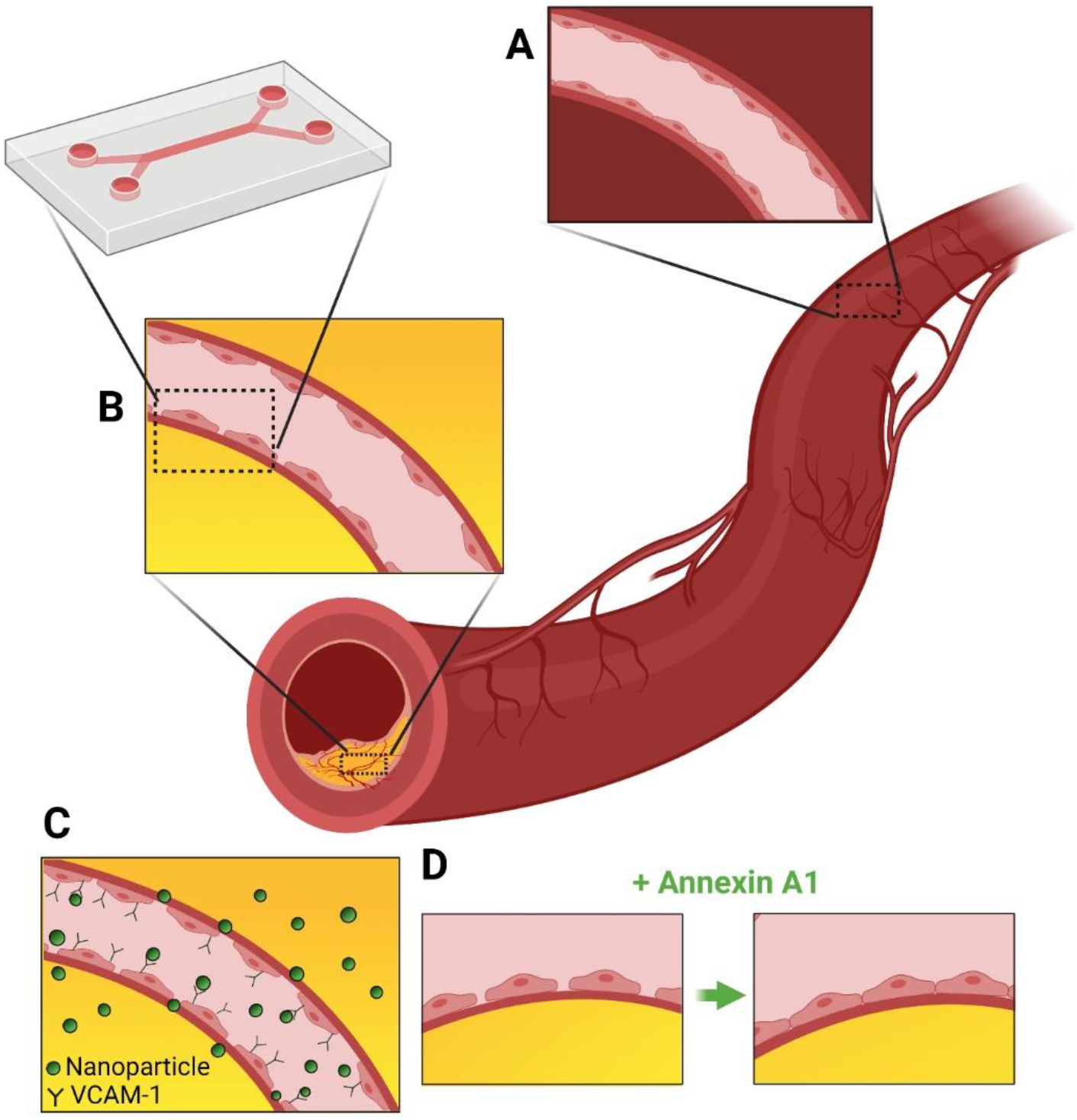
(A) Structure of a *vasa vasorum* on a healthy vessel wall. (B) Structure of a neovessel sprout from a *vasa vasorum* into an atherosclerotic plaque. The endothelium on these intraplaque neovessels is inflamed and Dys-En. We developed a Dys-En-a-chip in this study and used it to investigate: C) nanoparticle targeting to atherosclerotic lesions based on unique targeting opportunities provided by Dys-En (i.e. enhanced permeability and VCAM-1 overexpression), and D) restoration and normalization of Dys-En by an anti-inflammatory and resolving mediator (i.e. Annexin A1).

The importance of Dys-En in atherosclerotic therapy can be divided into two main categories: 1) targeting of drug delivery systems to atherosclerotic plaques based on unique targeting opportunities provided by Dys-En (**Figure 1C**) and 2) restoration and normalization of Dys-En to prevent chronic inflammation (**Figure 1D**). Targeted nanoparticles (NPs) are being intensively investigated for diagnosis and treatment of atherosclerosis ^16–19^, and Dys-En provides unique opportunities in order to target NPs to atherosclerotic plaques ^16^. Dys-En cannot regulate the barrier integrity, which leads to disrupted cell-cell junctions, thus enhancing permeability ^20,21^. Therefore, NPs can extravasate from the blood circulation into the plaque ^9,16^. Moreover, overexpression of adhesion molecules such as vascular cell adhesion molecule 1 (VCAM-1) is one of the key biological manifestation of a Dys-En ^22,23^. Both enhanced permeability and overexpression of VCAM-1 are promising strategies to target NPs to atherosclerotic plaques ^24–26^ (**Figure 1C**). Furthermore, Dys-En permits lipoproteins and leukocytes to enter the intima, which initiates chronic inflammation and contributes to plaque growth, especially in early atherosclerosis ^5,27^. Therefore, normalization and restoration of a compromised and leaky endothelium by novel anti-inflammatory and resolving mediators (e.g. Annexin A1) can be a potent therapeutic strategy for atherosclerosis ^28,29^ (**Figure 1D**).

In this study, we first fabricated a three-layered microchip, consisting of two polydimethylsiloxane (PDMS) layers separated by a semi-permeable membrane layer. Next, endothelial cells were cultured on the membrane using microfluidic cell culture techniques in order to develop a biomimetic model of endothelium-on-a-chip (En-on-a-chip). The established En-on-a-chip was inflamed using a pro-inflammatory cytokine to develop a dysfunctional-endothelium-on-a-chip (Dys-En-on-a-chip), which could successfully mimic the enhanced permeability, overexpression of VCAM-1, and the shear condition of a Dys-En. Then, we used the developed biomicrofluidic models to 1) screen and understand VCAM-1 targeted NP permeability and binding to Dys-En as a function of NP biophysicochemical properties and 2) investigate the restorative effect of Annexin A1 on Dys-En. A few studies have investigated binding of VCAM-1 targeted NPs to Dys-En under microfluidic shear conditions ^30–33^. However, a comprehensive study to investigate the effect of NP properties (i.e. size, VCAM-1 peptide density, zeta potential, and poly dispersity index (PDI)) on NP permeability and binding across healthy and Dys-En-on-a-chip is yet to be reported. Additionally, we chose to investigate the pro-resolving effects of Annexin-A1 on Dys-En because this protein has recently attracted a significant attention in the field of anti-inflammatory drug discovery ^29,34,35^. Annexin A1 can mimic how inflammation naturally subsides in the body, thus it has fewer side effects than current anti-inflammatory drugs ^29^. It has also been reported that Annexin A1 can show positive effects to restore the damaged blood-brain barrier (BBB) ^36–38^. However, the restorative and pro-resolving effects of Annexin A1 on Dys-En as a therapeutic strategy for CVD is yet to be investigated.

## 2. Results and Discussions

### 2.1. Development of En-on-a-chip and Dys-En-on-a-chip

In the first instance, we developed a microfluidic platform incorporating a monolayer of endothelial cells exposed to flow and shear stress, to be able to mimic the physiological environment of a vessel. For this purpose, we used a microfluidic chip consisting of three layers. The upper and lower layers were made of PDMS and fabricated by standard photolithography (**Figure S1**) and soft lithography procedures. Each PDMS layer contained a channel with the dimensions of 6 × 0.4 mm as the main channel, which was connected to two side channels with the dimensions of 4 × 0.4 mm. Two circles (2 mm diameter) were considered as the access points to the channels to prevent leakage at the inlet and outlet points. A thin layer of semi-permeable polyethylene terephthalate (PET) membrane was bonded between these PDMS layers. This membrane not only separated the upper and lower channels, but also provided a suitable support for culturing of endothelial cells. The schematic and photographic representation of the fabricated chip can be seen in **Figure 2A** and **Figure 2B-D**, respectively. After fabrication of the chip, the porous membrane was treated with fibronectin to promote the cell attachment. Human umbilical vein endothelial cells (HUVECs) were then cultured on the membrane by microfluidic cell culture techniques. The schematic representation of the developed biomicrofluidic system and the full coverage of the membrane by the cells can be seen in **Figure 2E** and **Figure 2F**, respectively. Various drugs (e.g. Annexin A1) and NPs (e.g. VCAM-1 targeted NPs) can be perfused through the upper channel and their interaction with the endothelium under physiological shear conditions can be studied not only by collecting the samples from the lower channel, but also by using microscopy techniques. The components of the microfluidic cell culture setup developed in this study are shown in **Figure S2A**. The system allows long-term culture of HUVECs under physiological flow conditions, inspection of the cells after seeding and during their growth, and accurate control of the flow rate, and thus shear stress. This setup was used to run the experiments for eight channels (i.e. four chips), simultaneously. A photographic representation of the setup can be seen in **Figure S2B**.

**Figure 2.**
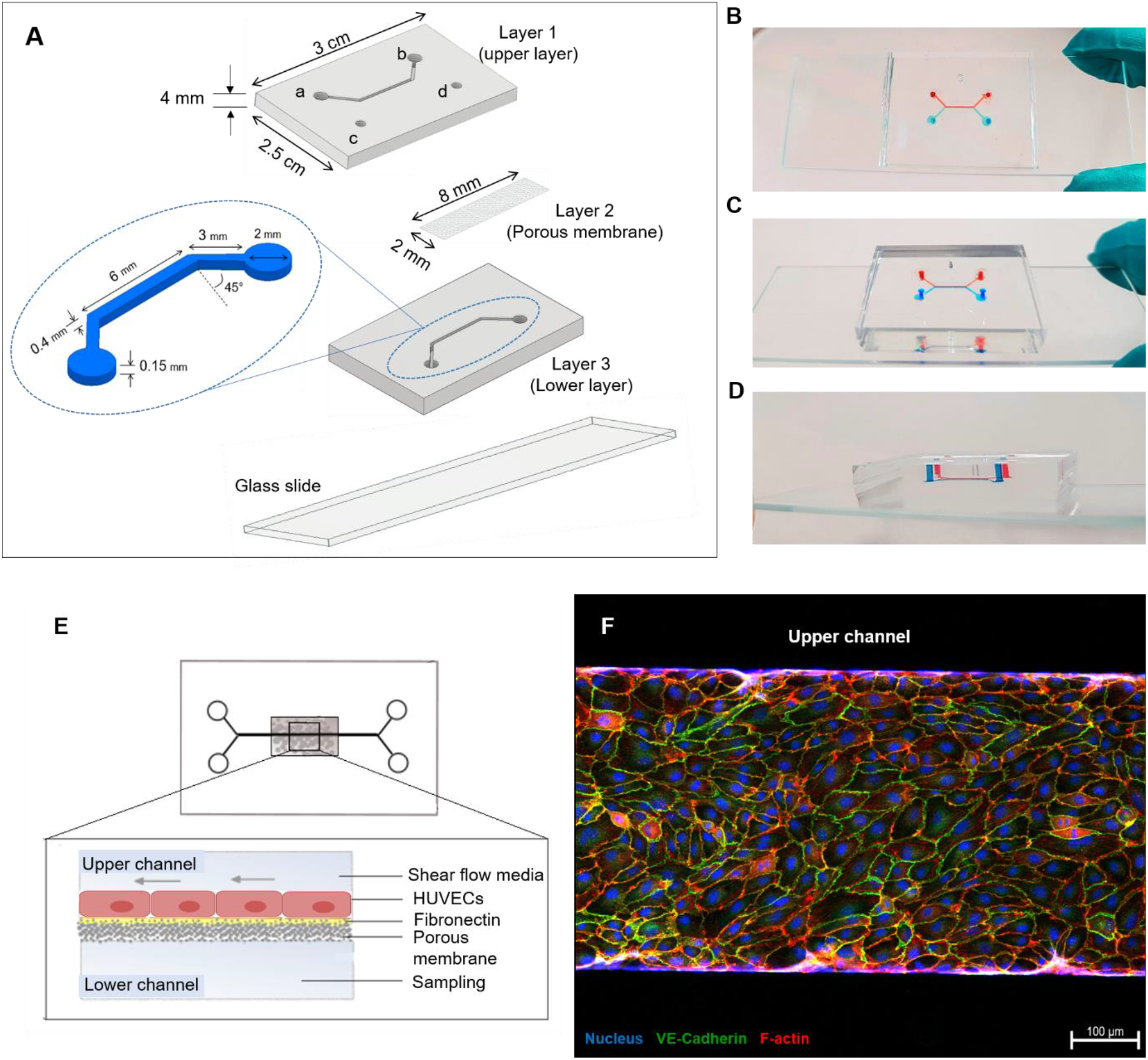
Development of the En-on-a-chip. (A) Schematic representation of the three-layered chip fabricated in this study and the dimensions of the chip and channels. In this figure, a and b show the inlet and outlet of the upper channel, c and d show the inlet and outlet of the lower channel. Photographic representation of the chip from (B) top view, (C) perspective view, and (D) side view. (E) Schematic representation of the chip after culturing the cells on the membrane. (F) Confocal image showing the top view of the upper channel proves the full coverage of the membrane by HUVEC cells (flow rate: 10 µL/min, scale bar: 100 µm). Cells were stained for nucleus (blue), VE-cadherin (green), and F-actin (red).

Based on in vivo measurements and fluid mechanical models, vessels prone to atherosclerosis and dysfunction show decreased shear stress (< 4-5 dyne/cm^2^) ^39,40^. Decreased shear stress can disrupt vascular wall functions (e.g. reduction in endothelium repair, decreased production of nitric oxide synthase (NOS), elevated production of reactive oxygen species (ROS), and higher permeability of lipoproteins and leukocytes) and therefore results in Dys-En and early-stage atherosclerosis ^40^. On the other hand, obstructive plaques in advanced atherosclerosis can significantly increase the shear stress ^41^. In this study, we aimed to focus on early-stage atherosclerosis. Therefore, we selected to use lower shear stress (< 4-5 dyne/cm^2^). The optimized shear stress for microfluidic cell-culture not only should be in line with reported in vivo conditions, but also should result in elongation of the cells along the direction of the flow and support a healthy cell culture without disruption of the cellular integrity. In order to find an optimized shear stress, we performed the microfluidic cell-culture with the shear stress of 0, 0.1, 1, 2, and 4 dyne/cm^2^. Based on Equation (1), the required flow rates for the shear stresses of 0, 0.1, 1, 2, and 4 dyne/cm^2^ were calculated to be 0, 1, 10, 20, and 40 µL/min, respectively.

As can be seen in **Figure 3**, the low shear stress (0 and 0.1 dyne/cm^2^) did not result in the complete elongation of the cells along the direction of the flow. On the other hand, high shear stress (2 and 4 dyne/cm^2^) disrupted cellular connections and did not support healthy cell culture and growth. Moderate shear stress (1 dyne/ cm^2^) showed elongation of the cells and complete barrier integrity. Therefore, we selected the shear stress of 1 dyne/cm^2^ (flow rate of 10 µL/min) as an optimized condition for HUVEC culture, and development of the En-on-a-chip and Dys-En-on-a-chip in this study.

**Figure 3.**
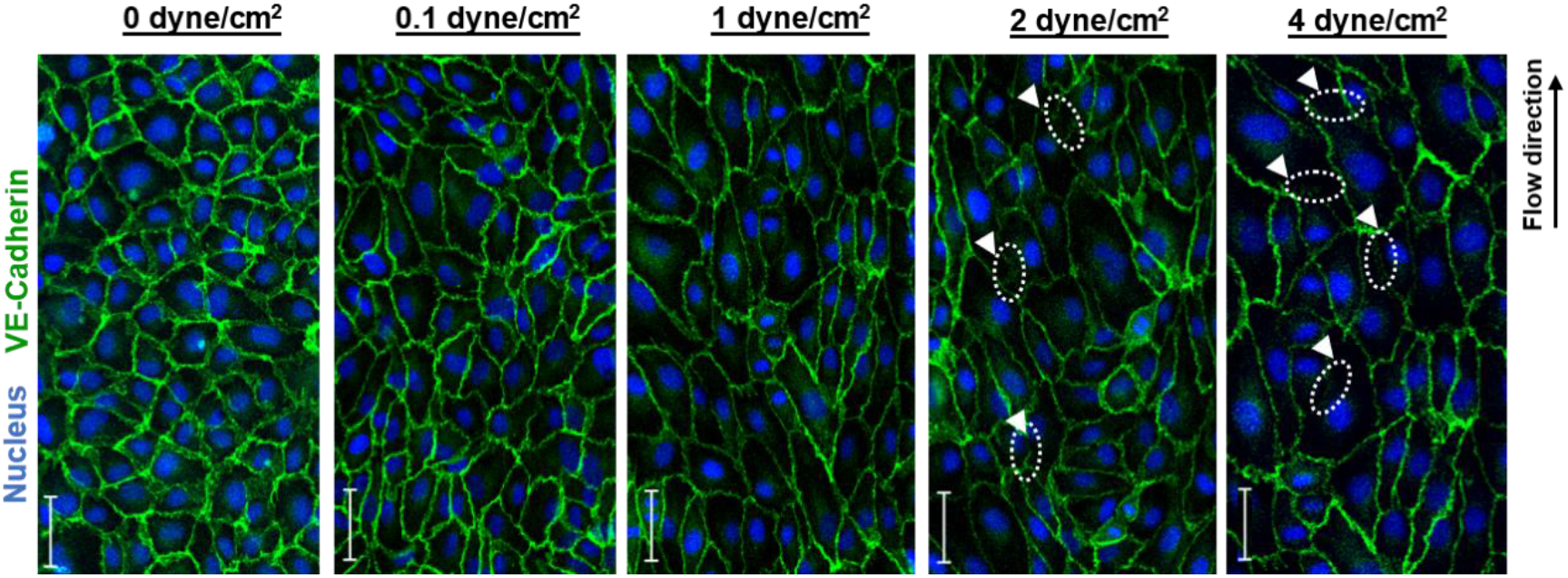
Confocal images of En-on-a-chip under different shear conditions (i.e. 0, 0.1, 1, 2, and 4 dyne/cm^2^). Cells were stained for VE-cadherin (green) and nucleus (blue) to visualize the orientation of the cells along the flow direction and endothelium integrity. Cells cultured with the shear stress of 0 and 0.1 dyne/cm^2^ did not show the complete elongation of the cells. Shear stress of 2 and 4 dyne/cm^2^ resulted in disrupted cellular connections and did not support healthy cell culture and growth. Some of the disrupted areas are marked by white arrowheads. Moderate shear stress (1 dyne/cm^2^) resulted in the elongation of the cells, while supporting the cellular integrity and selected as the optimal shear stress to culture HUVECs on chip in this study, (Scale bar: 50 µm).

After culturing of En-on-a-chip, tumor necrotic factor-α (TNF-α) was used to induce Dys-En. TNF-α is a proinflammatory cytokine, which is highly involved in the pathogenesis of cardiovascular disorders ^42,43^. TNF-α plasma levels in patients suffering from cardiovascular disorders was reported to be ~ 20 ng/ml ^44^. In this study, media containing 20 ng/ml of TNF-α was perfused through the developed En-on-a-chip and then three indicators of a Dys-En, including the loss of vascular endothelial cadherin (VE-cadherin), formation of actin stress fibers, and overexpression of VCAM-1 receptors, were investigated by immunostaining and fluorescent microscopy (**Figure 4**).

**Figure 4.**
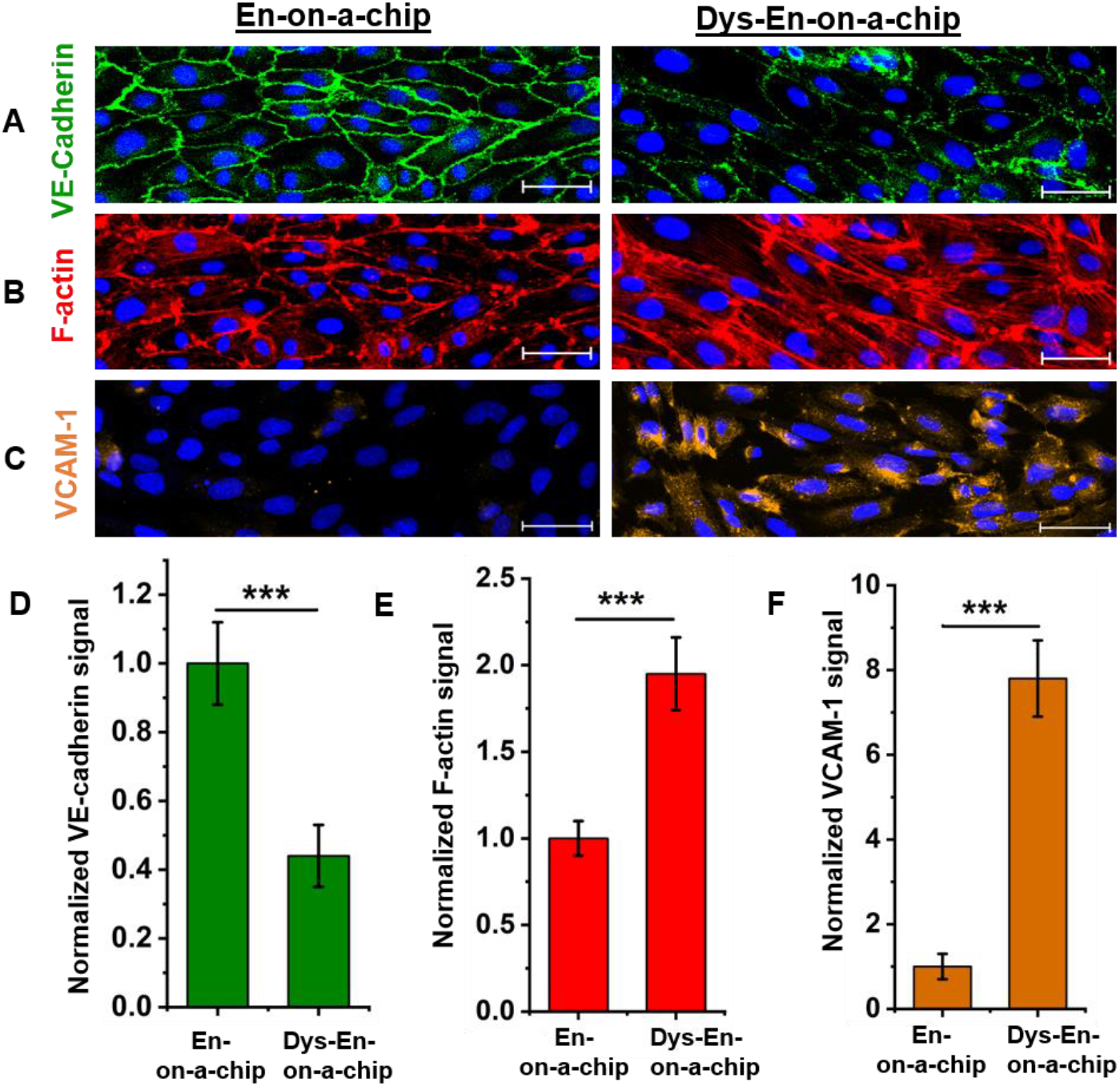
Confocal images of (A) VE-cadherin, (B) F-actin, and (C) VCAM-1 for En-on-a-chip and Dys-En-on-a-chip. In order to establish Dys-En, HUVECs were stimulated with TNF-α (20 ng/ml, 12 hours, 10 µl/min). The images are representative of 3 independent experiments (Scale bar: 50 µm). Quantification of (D) VE-cadherin, (E) F-actin, and (F) VCAM-1 signals confirmed the loss of VE-cadherin, formation of actin stress fibers, and overexpression of VCAM-1, which are known as hallmarks of a Dys-En. Data show mean ± SD (n=5-7) and *** represents statistically significant difference (p-value < 0.001).

VE-cadherin is a cell-cell adhesion protein, which plays a key role in endothelial barrier integrity and vascular permeability ^45^. The loss of VE-cadherin in endothelial cells results in endothelial barrier disruption and thus enhanced permeability ^21,46^. In this study, the established En-on-a-chip showed the assembly of VE-cadherin molecules at the cellular periphery, where they form tight borders between endothelial cells. On the other hand, Dys-En-on-a-chip showed patchy expression of VE-cadherin and loss of VE-cadherin connectivity at the cellular borders, which is an indicator of intercellular gap formation and enhanced permeability (**Figure 4A**). Furthermore, actin networks participate in endothelial barrier integrity by binding to junctional proteins and stabilizing them ^47^. Therefore, healthy En shows the arrangement of actin fibers at the cellular borders to establish endothelial junctions. In contrast, Dys-En shows the formation of actin fibers in central parts of the cells, known as actin stress fibers ^47^. These fibers cause transcellular tensions and cell contractions, and finally disrupt the barrier integrity ^47^. In this study, the developed En-on-a-chip showed the arrangement of actin fibers at the cellular periphery to form actin cortical rims. On the other hand, the developed Dys-En-on-a-chip could successfully show the formation of actin stress fibers, which were prolonged in the central domain of the endothelial cells (**Figure 4B**). Moreover, VCAM-1 immunostaining was performed to investigate the expression of this molecule on the developed En-on-a-chip and Dys-En-on-a-chip. VCAM-1 is an immunoglobulin-like adhesion molecule, which is normally absent on healthy endothelial cells but rapidly shows overexpression on inflamed and dysfunctional endothelial cells ^23,48^. The developed Dys-En-on-a-chip in this study, successfully showed a significant increase in VCAM-1 expression compared to the En-on-a-chip (**Figure 4C**). In order to quantify the fluorescent signals, imageJ software was used. The quantification of VE-cadherin, F-actin, and VCAM-1 signals respectively showed 2.2-fold decrease, 1.9-fold increase, and 7.8-fold increase in Dys-En-on-a-chip compared to En-on-a-chip. Therefore, the quantification results confirmed the loss of VE-cadherin, formation of actin stress fibers, and overexpression of VCAM-1 in the developed Dys-En-on-a-chip in this study (**Figure 4D-F**).

### 2.2. Permeability and Binding of VCAM-1 Targeted NPs across En-on-a-chip and Dys-En-on-a-chip

In order to gain understanding of which biophysicochemical properties of NPs (i.e. size, charge, PDI, and targeting peptide density) has the most influence on NP binding and translocation across the Dys-En under biomimetic microfluidic condition, a small library of VCAM-1 targeted polystyrene NPs was synthesized. To this end, amino-functionalized and rhodamine-labeled polystyrene NPs with the size of 30, 60, 120, and 250 nm were abbreviated as NP1, NP2, NP3, and NP4, respectively. The NPs were first PEGylated using maleimide polyethylene glycol *N*-hydroxysuccinimidyl ester (Mal-PEG-NHS) and methoxy polyethylene glycol *N*-hydroxysuccinimidyl ester (m-PEG-NHS) (**Figure 5A**). Subsequently, the PEGylated NPs were conjugated with a VCAM-1 targeting peptide (VHPKQHR-GGGC) (**Figure 5B**). After NP preparation, they were characterized for their size, zeta potential, PDI, and peptide density (**Figure 6A-D**), and then the permeability and binding of the NPs across En-on-a-chip and Dys-En-on-a-chip were studied (**Figure 6E-H**).

**Figure 5.**
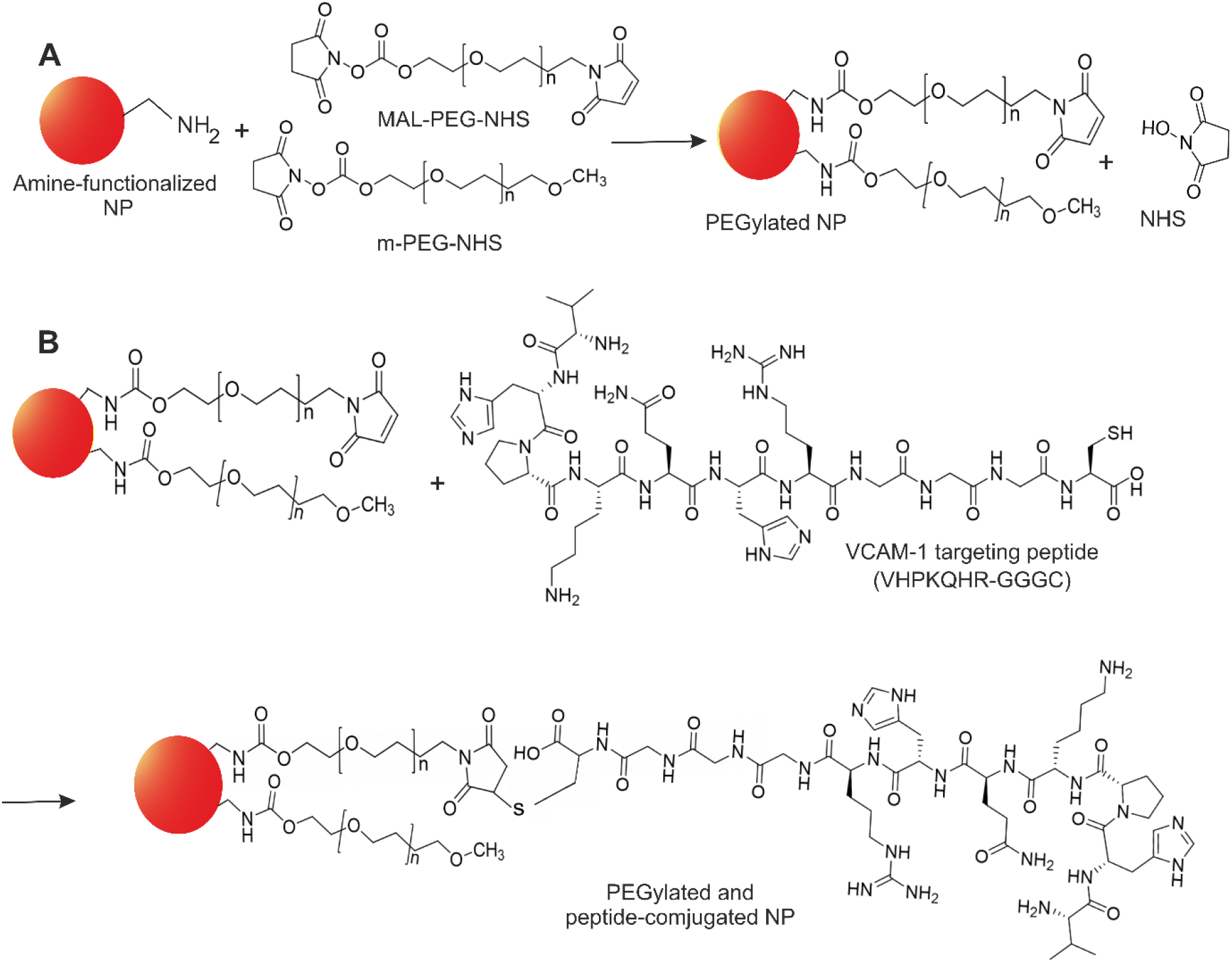
Development of VCAM-1 targeting NPs in this study. (A) PEGylation of the NPs using an amide bond coupling between NHS ester on the PEG polymers and the amino functionality on the surface of the polystyrene NPs. (B) The conjugation of the VCAM-1 targeting peptide to the surface of the NPs based on maleimide-thiol reaction between the maleimide groups on the surface of the NPs and the SH group of the peptide.

**Figure 6.**
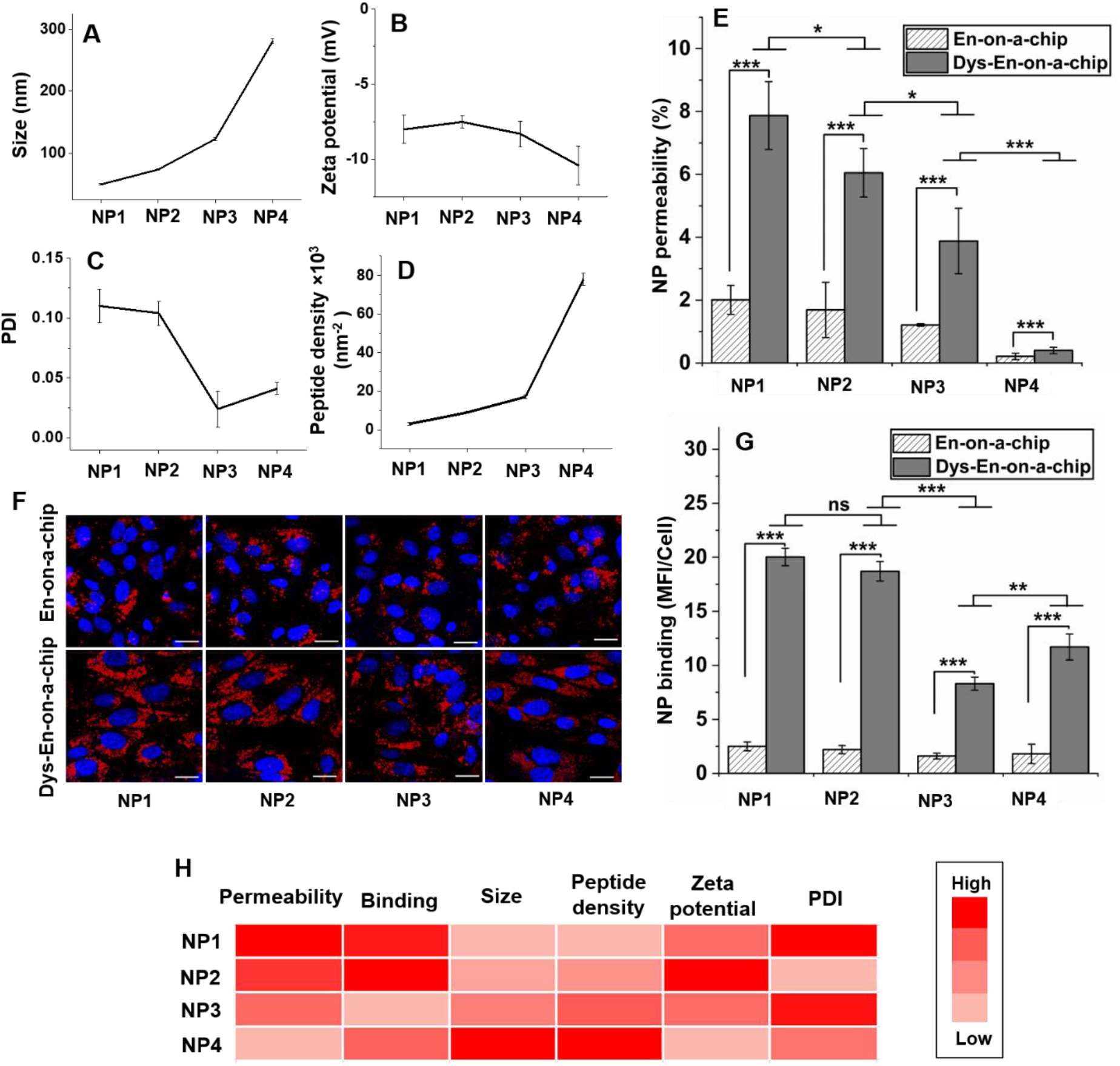
Characterization and screening of the NP library across En-on-a-chip and Dys-En-on-a-chip in this study. Characterization of the NPs for their (A) size, (B) zeta potential, (C) PDI, and (D) peptide density. (E) Permeability of the NPs across En-on-a-chip and Dys-En-on-a-chip. (F) Representative images showing the binding of the NPs to En-on-a-chip and Dys-En-on-a-chip. Red and blue colors show NPs and cell nuclei, respectively (scale bar: 20 µm). (G) Quantification of the binding results. (H) Heat-map summarizing the effect of NP properties on their permeability and binding across Dys-En-on-a-chip. The results suggest that NP size is the most dominant factor, and smaller NPs show higher permeability and binding. In this figure, data show mean ± SD of at least 3 independent experiments. *, **, *** represent statistically significant difference with p-value of <0.05, <0.01, and <0.001, respectively. ns represents non-significant differences.

The permeability of NP1, NP2, NP3, and NP4 across En-on-a-chip was 2.01 ± 0.46 %, 1.69 ± 0.88 %, 1.21 ± 0.04 %, and 0.21 ± 0.10 %, respectively. For Dys-En-on-a-chip, the permeability of all NPs was higher than their permeability across En-on-a-chip due to intercellular gaps, disrupted junctions, and compromised integrity of Dys-En. The permeability of NP1, NP2, NP3, and NP4 across Dys-En-on-a-chip was respectively 7.87 ± 1.08 %, 6.05 ± 0.77 %, 3.88 ± 1.04 %, and 0.51 ± 0.10 % (**Figure 6E**). The binding of the NPs to En-on-a-chip and Dys-En-on-a-chip was investigated by fluorescent microscopy. The representative images can be seen in **Figure 6F**. To quantify the results, mean fluorescent intensity (MFI) of rhodamine (i.e. MFI of the NPs) and the exact number of nuclei (i.e. the exact number of cells) were obtained and then MFI/cell was calculated. The binding of NP1, NP2, NP3, and NP4 to En-on-a-chip was respectively 2.5 ± 0.41, 2.2 ± 0.38, 1.6 ± 0.27, and 1.8 ± 0.9. However, their binding to Dys-En-on-a-chip was found to be 20.02 ± 0.8, 18.7 ± 0.9, 8.3 ± 0.6, and 11.7 ± 1.2 (**Figure 6G**).

Consequently, based on the above-mentioned data: 1) The permeability across Dys-En was in the following order: NP1 > NP2 > NP3 > NP4. So, the smaller the NPs are, the higher the permeability is. The larger NPs showed lower permeability, even though they had higher peptide density. These results suggest that size governs NP permeability regardless of peptide density. 2) NP binding to Dys-En was in the following order: NP1 and NP2 (no significant difference) > NP4 > NP3. So, the smaller NPs showed the highest binding even though they had the lowest peptide density. NP4, which had the highest peptide density among all groups, was in the next place, followed by NP3. Therefore, the results suggest that NP binding is firstly governed by size, then peptide density. 3) In this study, no trends were found between permeability and binding data with PDI and zeta potential. The summary of the results can be seen in **Figure 6H**, which shows the effect of the NP properties on their permeability and binding across Dys-En-on-a-chip.

Incorporating a porous membrane into a microfluidic device has been previously used to study the permeability of fluorescent tracers across endothelial layers ^49–52^. Although the results of these studies can be informative about the integrity and disruption of various endothelial layers, they cannot be used in the field of nanomedicine to design optimal NPs to traverse these barriers. The reason is that the hydrodynamic diameter of these tracers is much smaller than the diameter of typical NPs used in nanomedicine. Only a few studies have been reported on the permeability of NPs across healthy and/or Dys-En under microfluidic flow. For instance, Wang et al. studied the permeability of two types of NPs with different stiffnesses but similar sizes (~ 70 nm) across TNF-α stimulated HUVECs ^53^. The results showed a similar permeability for these NPs, which indicates the dominant effect of size on permeability. Also, Kim et al. studied the permeability of lipid NPs with a size of ~ 70 nm across non-stimulated and TNF-α stimulated HUVECs ^9^. The results showed a significant increase of NP permeability across TNF-α stimulated cells compared to non-stimulated cells, which is in line with our findings. However, none of these studies investigated the effect of NP properties such as size and peptide density on their permeability. Additionally, Sasaki et al. developed a mathematical model to investigate the relationship between the size of NPs and their permeability across a microporous membrane under microfluidic flow ^54^. Their model confirms the higher permeability of smaller NPs, but the main limitation of their study is that the membrane was without any cells. Therefore, the effect of NP-cell interactions on permeability was not considered.

The VCAM-1 binding to its ligand has been reported to show catch-bond characteristics ^30,55^. The lifetime of a catch-bond initially increases by force because the molecules deform and lock more tightly, and then decreases like in a normal bond ^56^. Therefore, any mechanical force like shear stress can affect VCAM-1 binding. Despite the important role of shear in this field, only a few studies have used shear generating systems when studying the effect of NP properties on VCAM-1 binding ^30,49,57^. For instance, Kusunose et al. synthesized a VCAM-1 targeting liposome with a size of ~ 100 nm and studied the effect of peptide density on VCAM-1 binding under microfluidic flow. The results showed that the NP binding significantly increases by increasing the peptide density ^30^. Furthermore, Campos et al. investigated the effect of NP shape (rod-like and spherical NPs with a size of ~ 200 nm) on VCAM-1 binding under microfluidic flow. The results revealed that the binding of rod-like NPs was significantly higher than the binding of spherical NPs ^57^. However, all studies were based on single-sized NPs, and the effect of NP size on VCAM-1 binding has not been reported to our knowledge.

### 2.3. Restorative Effect of Annexin A1 on Dys-En-on-a-chip

To study the effect of Annexin A1 on Dys-En, we first measured the translocation of Lucifer Yellow, as an endothelial permeability marker, across the developed En-on-a-chip, Dys-En-on-a-chip, and Annexin A1 treated Dys-En-on-a-chip (**Figure 7A**). Lucifer Yellow is a hydrophilic and fluorescent tracer molecule, with wide stokes shift and distinct excitation/emission wavelength, making it an ideal marker for robust permeability studies in cell models ^58^. The average intensity of the translocated Lucifer Yellow across Dys-En-on-a-chip was 3.5-fold higher than En-on-a-chip, which indicates the enhanced permeability and compromised integrity of the developed Dys-En compared to the healthy En. However, Annexin A1 treatment could significantly lower the Lucifer Yellow translocation across the treated Dys-En. The average intensity of translocated Lucifer Yellow across Annexin A1 treated Dys-En-on-a-chip was 1.8-fold lower than non-treated Dys-En-on-a-chip, which confirmed the endothelial tightening and the rescuing effect of Annexin A1 on permeability of Dys-En (**Figure 7B**).

**Figure 7.**
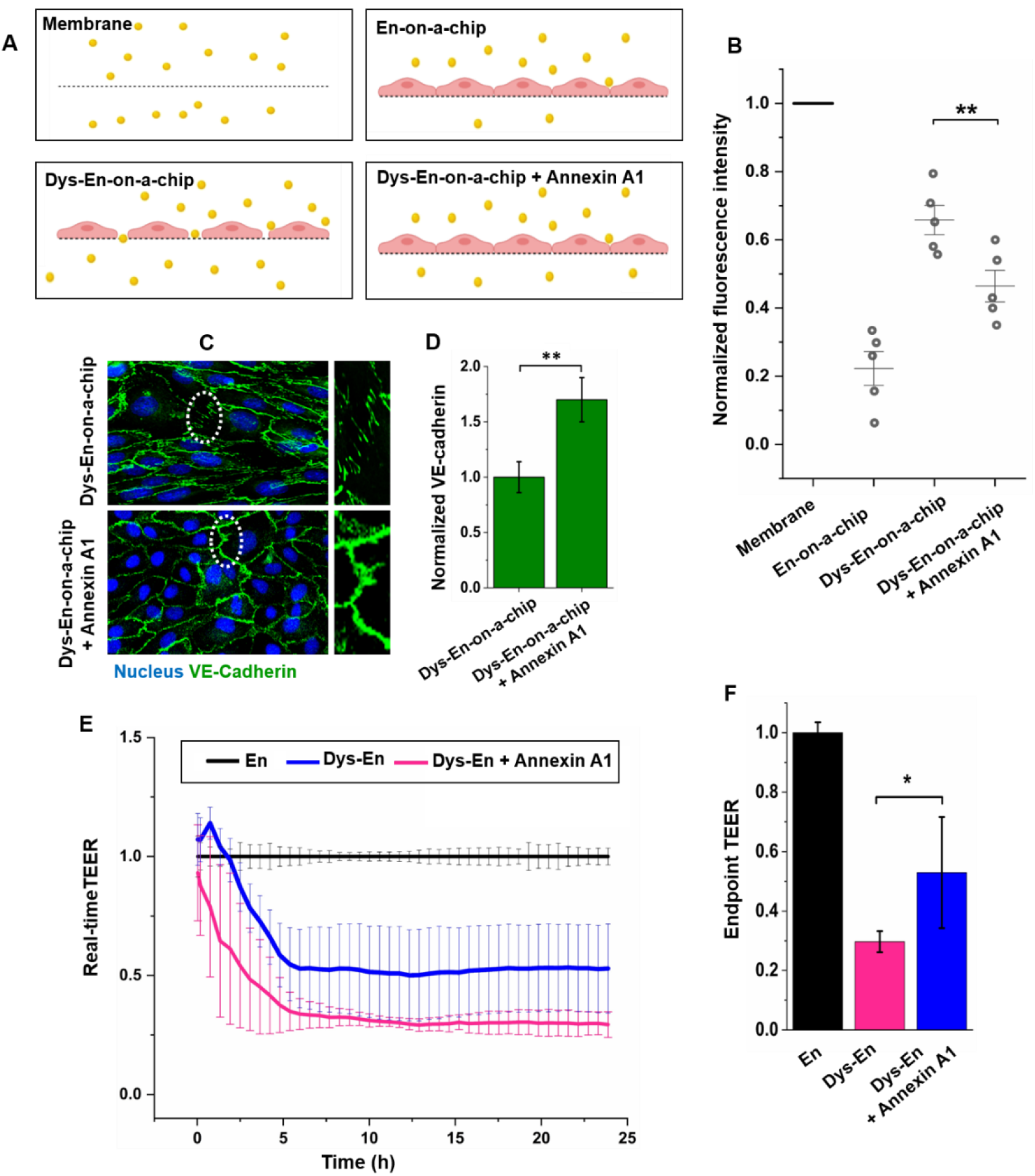
Restorative effect of Annexin A1 (20 µg/ml) on Dys-En. (A) Schematic representation of Lucifer Yellow permeability across a membrane (without any cells), En, Dys-En, and Annexin A1 treated Dys-En. (B) Lucifer Yellow permeability results (5 µg/ml, 30 min, 5 µl/min). Annexin A1 treatment decreased Lucifer Yellow permeability, confirming the endothelial tightening. (C) Immunostaining and confocal microscopy of VE-cadherin. White frames show regions of interests (ROIs) enlarged in the second panel. (D) Quantification of VE-cadherin signals by ImageJ. Annexin A1 treatment increased the expression of this critical component of endothelial junctions. (E) Real-time and (F) endpoint TEER measurements. Annexin A1 treatment increased the TEER values, confirming its positive effect on endothelial integrity. In this figure, data show mean ± SD (n=3-5). * and ** represent statistically significant difference with p-value of <0.05 and <0.01, respectively.

Furthermore, immunostaining of VE-cadherin was performed to investigate the distribution of this adherens junction component for Dys-En-on-a-chip before and after Annexin A1 treatment (**Figure 7C-D**). The results of confocal microscopy displayed a patchy distribution of VE-cadherin and loss of cell-cell connectivity in non-treated Dys-En-on-a-chip. However, the treated Dys-En-on-a-chip showed more distribution of VE-cadherin molecules at the cellular borders and higher connectivity of the junctions (**Figure 7C**). The quantification of fluorescent signals by ImageJ confirmed a ~ 1.7-fold increase of VE-cadherin signals in treated Dys-En-on-a-chip, compared to non-treated Dys-En-on-a-chip (**Figure 7D**). Therefore, Annexin A1 treatment of the developed Dys-En-on-a-chip could inhibit the loss of VE-cadherin and/or enhance the expression of this critical component of adherens junctions, further confirming the potential use of Annexin A1 as a valuable drug candidate for cardiovascular disease.

Additionally, we measured transendothelial electrical resistance (TEER) for healthy En, Dys-En, and Annexin A1 treated Dys-En, using an automated multi-well CellZscope device. TEER measurement is a non-destructive and label-free method to study barrier integrity. Intact and tightly connected endothelial cells are resistant to the passage of the electrical current but disrupted endothelial layers allow the passage of the electrical current more easily. Therefore, there is a direct relationship between the measured TEER and the integrity of the endothelial layer ^59^. **Figure 7E** shows the real-time TEER, and **Figure 7F** shows the measured TEER at the end of the experiment (after 24 hours), termed as endpoint TEER. As shown in these figures, Annexin A1 showed a positive effect on integrity of endothelial layers. The endpoint TEER for Dys-En was 3-fold less than healthy En, which confirms the disrupted barrier integrity of the developed Dys-En model. However, Annexin A1 treatment could increase the TEER values, and the endpoint TEER for treated Dys-En was 1.5-fold higher than non-treated Dys-En, which indicates the restorative effect of this protein on endothelial integrity.

The above-mentioned results (i.e. Lucifer Yellow permeability, VE-cadherin microscopy, and TEER measurements) confirmed the positive effect of Annexin A1 on Dys-En. We speculate that the effect of Annexin A1 to restore Dys-En is due to the inhibition of RhoA GTPase pathway, also known as the RhoA-ROCK pathway ^37,38^. Future studies could address this probability in order to gain further mechanistic insight into Annexin A1 effects on Dys-En.

## 3. Conclusion

In this study, a microchip consisting of two parallel channels separated by a semi-permeable membrane, which is a simple but robust design for on-chip permeability studies, was utilized to establish a Dys-En-on-a-chip model. The developed biomicrofluidic model not only can mimic the relevant pathological shear stress but also showed loss of adherens-junctions at cellular borders, change of cytoskeleton organization to form actin stress fibers, and upregulation of VCAM-1 on endothelial cells, which are all indicators of a successful Dys-En biomimetic *in vitro* model. The developed platform was used to screen a library of VCAM-1 binding NPs with different properties. Based on the NP translocation assessments and microscopy studies, smaller NPs showed higher permeability and binding, and NP size dominated the NP targeting to Dys-En. Moreover, the restorative effect of Annexin A1 on Dys-En was investigated. Treatment of Dys-En-on-a-chip with Annexin A1 resulted in restoration of adherens-junctions and enhancement of the barrier integrity, which confirms the potential application for this drug to be used clinically for the treatment of CVD such as atherosclerosis.

## 4. Experimental Section

### Materials

SU-8 2075 and PDMS (Sylgard 184) were purchased from MicroChem (USA) and Dow Corning (USA), respectively. (3-aminopropyl) triethoxysilane (APTES), Perfluorodecyltrichlorosilane (FDTS), phosphate buffer saline (PBS), fetal bovine serum (FBS), penicillin-streptomycin, fibronectin from human plasma, trypsin, ethylene diamine tetraacetic acid (EDTA), Triton X-100, bovine serum albumin (BSA), dimethyl sulfoxide (DMSO), cysteine, Lucifer Yellow, and ethanol were provided by Sigma-Aldrich (Denmark). HUVECs (PCS-100-013), vascular cell basal medium (PCS-100-030), and vascular endothelial growth factor (VEGF) kit (PCS-100-041) were obtained from American Type Cell Culture (ATCC, USA). Transwell inserts, containing a PET porous membrane with the pore size of 1 µm were purchased from Corning (UK). TNF-α and recombinant human Annexin A1 were obtained from R and D systems (USA). Alexa fluor488 VE-cadherin antibody was provided by Santa Cruz Biotechnology (USA). Alexa fluor647 Phalloidin, mouse monoclonal IgG1 anti-VCAM-1 antibody, and Alexa fluor555-conjugated goat anti-mouse IgG1 antibody were obtained from Invitrogen (Denmark). Polyether ether ketone (PEEK) tubing, Tygon tubing, PEEK micro-tight adaptor, paraformaldehyde solution, and Hoechst 33342 were obtained from Thermo Fisher Scientific (Denmark). Polystyrene NPs, which were functionalized by amine groups and labeled by rhodamine B, were provided in four different sizes (i.e. 30, 60, 120, and 250 nm) by Micromod (Germany). VCAM-1 targeting peptide (VHPKQHR-GGGC) was obtained from Mimotopes (UK). MAL-PEG-NHS (2 kDa) and m-PEG-NHS (2 kDa) were obtained from Nanocs (USA).

### Photolithography and Soft Lithography

Microfluidic channels were designed using CleWin software (PhoeniX Technologies, USA) and then patterned on a silicon wafer using SU-8 and standard photolithography procedure (see supporting information and **Figure S1**). Subsequently, silanization was performed to prevent the molds from sticking to PDMS. For this purpose, the molds were silanized by FDTS using a molecular vapor deposition system (MVD 100, Applied Microstructures, UK) for 60 min. After mold fabrication and silanization, PDMS soft lithography was performed ^60^. Briefly, PDMS and its curing agent were mixed (10:1 w/w ratio), degassed, poured over the mold, and baked at 80°C for 2 h. The cured PDMS slab was peeled from the mold and cut into single devices (i.e. upper and lower layers of the chip). The access holes to the channels were punched using a 1.25 mm biopsy puncher on the upper layer.

### Bonding of PDMS Layers and Membrane

The upper layer was bonded with a microporous PET membrane with the pore size of 1 µm. For this purpose, the protocol developed by Aran et al. was used ^61^. Briefly, the membrane was activated in an oxygen plasma chamber (600 mTorr, 100 W, Harrick Plasma, USA) for 1 min and then immersed in 5% APTES solution at 80°C for 20 min. Subsequently, the PDMS upper layer was activated in an oxygen plasma chamber for 20 s, immediately brought into contact with the treated membrane, and then heated at 80°C for 10 min. Finally, the bonded laminate and the PDMS lower layer were exposed to the oxygen plasma for 20 s and bonded together considering the alignment of the channels. For accurate alignment of the channels, a Dino-Lite microscope (AM7915MZTL, Dino-Lite, UK) was used. Lastly, the fabricated three-layered chip was placed in an oven (60°C, overnight) to assure the complete bonding of the layers.

### Cell Line and Cell Maintenance

HUVECs were cultured in vascular cell basal medium, which was supplemented with VEGF kit, 10% FBS, and 1% penicillin-streptomycin. The cells were cultured at 37°C in a humidified incubator with 5% CO_2_. The media was changed every second day. The cells were trypsinized after reaching 80% confluency, centrifuged at 100 g for 5 min, and resuspended in culture media at the required concentration. For all experiments, the cells were used between passages 3 to 7.

### Microfluidic Cell-culture Setup

The microfluidic cell-culture setup used in this study consists of a microfluidic flow system, a microscope, a chip holder, tubing, connectors, and reservoirs. The microfluidic flow system consists of a microfluidic pressure pump (MFCS-EX, with 8 channels, Fluigent, Germany), a flow sensor (FRP8, Fluigent, Germany), caps for pressurization of the media and collection reservoirs (Fluiwell-1C-15ml, Fluigent, Germany), and a software to control and monitor the flow rate (A-i-O software, Fluigent, Germany). The pressure pump was connected to a local compressed air tap, which was equipped by a high precision regulator for adjusting the suitable inlet pressure to run the pump (1300 mBar). The light microscope and a flexible tabletop microscope stand were purchased from Dino-Lite Inc. (AM7915MZTL microscope, and RK-04F stand, Dino-Lite, UK) in order to visualize the cells after cell-seeding and during culture. The chip holder with the total length, width, and height of 15 × 10 × 2 cm was fabricated using polymethyl methacrylate (PMMA) and a laser cutter machine (Epilog laser, USA). The fabricated chip holder contains four separate holders with the dimensions of 7.5 × 2.5 × 1 cm for four chips, which were bonded to microscopy slides. The tubing connected to the flow sensor was PEEK tubing (with the outer dimension of 0.75 mm and the inner dimension of 125 µm). The tubing connected to the chips was Tygon tubing (with the outer dimension of 1.5 mm and the inner dimension of 0.5 mm). These two tubes were connected to each other using a PEEK micro-tight adaptor (For schematic and photographic figures of the microfluidic setup, see supporting information and **Figure S2)**.

### Development of En-on-a-chip

Firstly, all connectors, fittings, tubing, filters, reservoirs, and caps were sterilized by autoclaving. Then, they were connected to the chips and the flow system inside a laminar flow hood under sterilized conditions. To ensure the sterilization of the chips and all components, the entire system was sterilized by perfusing 70% ethanol for 30 min at the flow rate of 20 µl/min. Subsequently, PBS was perfused for 30 min at the same flow rate. After preparation and sterilization of the setup, it was placed in a cell-culture incubator (37°C, 5% CO_2_) for the rest of the experiment duration. Then, 100 µg/mL fibronectin solution was perfused through the upper channels of the chips with the flow rate of 100 µl/min for 2 min, to coat the membranes. Then, the system was statically incubated with the perfused fibronectin for 1 h. Following this, HUVECs (1 x 10^7^ cells/ml) were perfused through the upper channels of the chips. The cell distribution was inspected using the microscope to ensure complete and even seeding of the cells on the membrane. The cells were incubated for 3 h under static condition to settle and adhere to the membrane. Subsequently, the continuous flow of the culture media was started at the flow rate of 10 µl/min (unless otherwise indicated). The condition of the cells was regularly inspected using the microscope of the system. After 48 h, the cells were found to reach confluency and establish a healthy En-on-a-chip.

### Development of Dys-En-on-a-chip

HUVECs were grown to confluency as mentioned above. Subsequently, TNF-α (20 ng/ml in culture media) was perfused through the upper channel for 12 h.

### Shear Stress Calculation

To calculate shear stress from the flow data in a microchannel with a rectangular cross-section, Equation (1) was used ^62,63^:

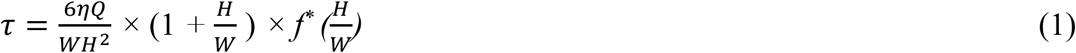

In this equation, *τ* and *Q* are shear stress and volumetric flow rate, respectively. *η* is the viscosity of the fluid and assumed to be equal to the viscosity of water at 37°C, which is ~10^−3^ Pa.s. *W* is the width and *H* is the height of the microchannel. 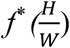 is a correction factor, which depends on the channel dimensions and the aspect ratio ^63^. For the designed microchannel in this study, the height-to-width ratio 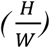 is 0.375. The calculated *f*^*\**^ for the channel with this aspect ratio is reported to be 0.7 ^63^.

### Immunostaining and Quantification

For all steps of immunostaining experiments, the reagent solutions were perfused through the upper channel at the flow rate of 100 µL/min for 2 min to deliver the reagents to the cells. Then, the cells were incubated with the reagents under a stopped-flow condition at room temperature. The solutions were introduced to the cells in this order: PBS to rinse away the culture medium, 4% paraformaldehyde solution for 20 min to fix the cells, PBS to rinse, 0.1% Triton X-100 for 30 min to permeabilize the cells, PBS to rinse, and 1% BSA solution for 1 h to block nonspecific bindings. Subsequently, cells were labeled with an immunostaining antibody. For VE-cadherin staining, cells were incubated with Alexa fluor488 VE-cadherin antibody (diluted 1:1000 in PBS) for 40 min. In the case of F-actin staining, cells were incubated with Alexa fluor647 Phalloidin (diluted 1:40 in PBS) for 1 h. For VCAM-1 staining, cells were first incubated with mouse monoclonal IgG1 anti-VCAM-1 antibody (diluted 1:50 in PBS containing 1% BSA) for 1 h as the primary antibody. Following that, cells were rinsed with PBS and incubated with Alexa fluor555-conjugated goat anti-mouse IgG1 antibody (diluted 1:1000 in PBS) for 1 h as the secondary antibody. For all experiments, nuclei staining was performed using Hoechst (diluted 1:2000 in PBS) for 30 min. Lastly, confocal images were captured using a laser scanning confocal microscope (LSM780, Zeiss, Germany).

In order to quantify the signals, three independent experiments were performed, 5-7 images of each sample were obtained, and then all images were quantified by imageJ software (ImageJ 1.47, USA). To quantify the VE-cadherin signals in an image, the threshold value was selected by the software and applied to the images. Then, the fluorescent intensity above the threshold value was calculated by the software, normalized to the number of nuclei in the image, and expressed as average VE-cadherin signal/cell. Using a single threshold value for all samples within an experiment, the average VE-cadherin signal/cell was calculated for En-on-a-chip and Dys-En-a-chip. Lastly, data obtained from Dys-En-a-chip were normalized to the data obtained from En-on-a-chip. To quantify F-actin and VCAM-1 signals, a similar quantification method was performed.

### Preparation of VCAM-1 Targeted Polystyrene NPs

For this purpose, polystyrene NPs, which were functionalized by amine groups and labeled by rhodamine B, were purchased from Micromod (Germany). NP solutions were first washed twice in phosphate buffer (20 mM, pH8). For this purpose, NPs were centrifuged at 10000 rpm, 4°C (30 min for NP1, NP2, and NP3, and 15 min for NP4) using Amicon ultra-0.5 centrifugal filter units (10 kDa cutoff, Millipore, UK) and then dissolved in 200 µL phosphate buffer. Then, NPs were PEGylated by a mixture of MAL-PEG-NHS and m-PEG-NHS. In this study, we used MAL-PEG-NHS and m-PEG-NHS with a molar ratio of 1:5 in order to avoid remaining of MAL groups on the surface of the NPs, which can react with thiol groups on cells and cell culture proteins. The PEG linkers were dissolved in DMSO, and then mixed with NP solutions at room temperature overnight. The molar ratio of amine groups to PEG in these reactions was 1:3. Next, the VCAM-1 targeting peptide was conjugated to the surface of the NPs. For this purpose, the PEGylated NPs were washed twice in PBS buffer containing 1mM EDTA, and then reacted with an excess amount of the peptide overnight at room temperature. Then, unreacted maleimide groups were quenched using 20mM cysteine solution. Lastly, the NPs were washed twice in PBS using Amicon centrifugal filter units as mentioned above.

### Characterization of VCAM-1 Targeted Polystyrene NPs

Size, zeta potential, and PDI of the NPs were measured by dynamic light scattering technique (DLS, Nano-ZS, Malvern, UK). In order to quantify the peptide density on the surface of the NPs (number of peptides per nm^2^ of NP surface or *#*_*peptide*_*/nm*^*2*^), the UV absorbance of the NPs was measured at 205 nm before and after peptide coupling using a spectrophotometer (Nanodrop 2000, Thermo Fisher Scientific, UK) and the difference of the absorbance (*ΔA*_*205*_) was obtained. Moreover, the extinction coefficient (*ε*_*205*_) of the peptide at 205 nm was calculated based on a model reported by Anthis et al. ^64^ and their online computation program (available at: http://spin.niddk.nih.gov/clore). Based on this model, the extinction coefficient of VHPKQHR-GGGC at 205 nm, is 40640 M^−1^ cm^−1^. Therefore, the concentration of the peptide was calculated based on *ε*_*205*_ and *ΔA*_*205*_ results, using the Beer-lambert law. Lastly, we calculated *#*_*peptide*_*/nm*^*2*^ by knowing the concentration of the peptide, NP size, and NP density (1.03 g/cm^3^).

### Permeability of VCAM-1 Targeted NPs across En-on-a-chip and Dys-En-on-a-chip

NP solutions (100 µg/ml in culture medium) was perfused through the upper channel at the flow rate of 5 µL/min. Moreover, the culture medium without any NPs was perfused through the lower channel at the flow rate of 1 µL/min. After 6 h, the solution containing translocated NPs was collected from the collection reservoir of the lower channel. This solution was analyzed using a fluorescence plate reader (Spark, Tecan Life Science, Switzerland) at the excitation/emission wavelength of 552/580 nm to measure the fluorescent intensity of the translocated NPs. Furthermore, standard curves were prepared to indicate the relationship between the intensity and NP concentration. Based on the intensity results and the standard curves, the concentration of the translocated NPs was calculated. Lastly, the concentration of the translocated NPs was divided by the initial concentration of the NPs to obtain % permeability.

### Binding of VCAM-1 Targeted NPs to En-on-a-chip and Dys-En-on-a-chip

For this purpose, the NP solution at the concentration of 100 µg/ml was perfused through the upper channel at the flow rate of 5 µL/min for 2 h. Subsequently, the cells were washed extensively to remove any unbound NP, fixed by 4% paraformaldehyde solution for 20 min, stained for nucleus by Hoechst (diluted 1:2000 in PBS) for 30 min, and then imaged using a laser scanning confocal microscope (LSM780, Zeiss, Germany). Zen Intellesis software (Zeiss, Germany) was used to quantify the images and obtain the MFI of NPs and the exact number of cells for each image. Then, MFI/cell was calculated.

### Modulation of Dys-En-on-a-chip by Annexin A1

Firstly, Dys-En-on-a-chip was developed as mentioned above. Then, 20 µg/ml recombinant human Annexin A1 was added to the TNF-α containing medium and perfused through the upper channel at the flow rate of 10 µl/min for 12 h.

### Lucifer Yellow Permeability Assay

Lucifer Yellow solution (5 µg/ml in culture medium) was perfused through the upper channel. The culture medium without any fluorescent tracer was perfused through the lower channel. The flow rate for both channels was set at 5 µl/min. After 30 min, the samples were collected from the collection reservoir of the lower channel. The fluorescence intensity was measured using a fluorescence plate reader (Spark, Tecan Life Science, Switzerland) with excitation/emission wavelength of 428/536 nm. For comparison, data were normalized to the fluorescence intensity of the translocated Lucifer Yellow across a bare membrane (a chip without any cells).

### TEER Measurements

TEER was measured using CellZscope (NanoAnalytics, Germany). Firstly, cells were cultured on Transwell inserts, containing a PET porous membrane with pore size of 1 µm. For this purpose, the membranes were treated with 100 µg/ml fibronectin for 1 h at 37°C. Then, HUVECs at the cell density of 0.2 × 10^6^ cell/inserts were seeded on the membranes and incubated for 24 h to form confluent endothelial layers (En group). Then, the media in the inserts was exchanged with the culture media containing 20 ng/ml TNF-α, and the inserts were incubated for 12 h (Dys-En group). Subsequently, 20 µg/ml Annexin A1 was added to the inserts (Annexin A1 treated Dys-En group). Lastly, 3-5 inserts of each group were placed inside the CellZscope device and the TEER was measured every 30 min, for the duration of 24 h. The data obtained from the Dys-En group and Annexin A1 treated Dys-En group were normalized to the data obtained from the En group.

### Statistical Analysis

OriginPro 10 (OriginLab, USA) was used to perform statistical analyses. All data were derived from at least 3 independent experiments and reported as mean ± SD. The comparison between groups were made by two-sample t-test. The p-values of <0.05, <0.01, and <0.001 were shown as *, **, and ***, respectively. The symbol ‘ns’ shows non-significant differences.

## Supporting information

Investigating VCAM-1 Targeted Nanoparticles and Annexin A1 Therapy using Dysfunctional-endothelium-on-a-chip

## Acknowledgement

This research was supported by a Lundbeck Fellowship awarded to N.K (R215-2015-4190).

## Conflict of Interest

The authors declare no conflict of interest.

## Notes

### Competing Interest Statement

The authors have declared no competing interest.

