## Supplementary material for "Investigating VCAM-1 Targeted Nanoparticles and Annexin A1 Therapy using Dysfunctional-endothelium-on-a-chip": Investigating VCAM-1 Targeted Nanoparticles and Annexin A1 Therapy using Dysfunctional-endothelium-on-a-chip

**Photolithography and Fabrication of SU-8 Master Molds.** For this purpose, 6-inch silicon wafers were first heated overnight in a 250 °C oven to remove moisture. Then, SU-8 2075 (MicroChem, USA) was spin-coated (RCD8 T spin coater, Süss MicroTek, Germany) on the silicon wafers to get a uniform layer of SU-8 on the surface of the wafers. The speed and duration of the spin-coating process to achieve the final thickness of 150 μm were adjusted based on our preliminary studies. The spreading speed and duration were 300 rpm and 30 s, followed by the spinning speed of 600 rpm for 60 s. Subsequently, soft bake (i.e. post-coating bake) was performed to evaporate the solvent and densify the coated layer. For this purpose, the wafers were placed on a programmable hot-plate, the temperature was increased from room temperature to 65 °C with the increasing rate of 5 °C/min, the wafers were baked at 65 °C for 10 min, the temperature was increased from 65 °C to 95 °C using the increasing rate of 5 °C/min, the wafers were baked at 95 °C for 45 min, and finally placed at room temperature to cool down. The reason for the temperature ramps during the soft-backing procedure was reduction of internal mechanical stresses. After the soft-backing step, the wafers and the photomask were placed in an aligner (Mask aligner MA6-2, Süss MicroTek, Germany). Then, UV light with the wavelength of 365 nm and exposure dose of 390 mJ/cm$^2$ was exposed to the wafer through the photomask, which contains the predefined patterns of the channels, to initiate the photopolymerization process. Therefore, the SU-8 layer started to crosslink in the defined sites. Subsequently, post-exposure bake was used to increase the diffusion rate of photoactive compounds and complete the photo-crosslinking procedure. The temperatures, durations, and temperature ramps were same as the soft-backing step. The wafers were then submerged in a SU-8 development bath, containing propylene glycol monoethyl ether acetate (MGMES, also known as mr-Dev 600, Sigma-Aldrich, Denmark) for 30 minutes to dissolve the non-crosslinked parts of the SU-8 layer and get the final patterns of the channels. Lastly, the wafers were cleaned by isopropyl alcohol (Sigma-Aldrich, Denmark) and dried.

To inspect the patterns on the fabricated SU-8 master molds, optical microscopy (Leica Microsystems, Germany) and profilometer (Dektak XTA stylus profilometer, Bruker, USA) were used. The microscopy results confirmed that the width of the channels was 400 μm. Moreover, based on the results from profilometer, the height of the channels was 150 μm. These results show that the fabrication process of the master molds was successfully performed. **Figure S1** shows the fabricated SU-8 mold in this study, the patterns, and the characterizations.

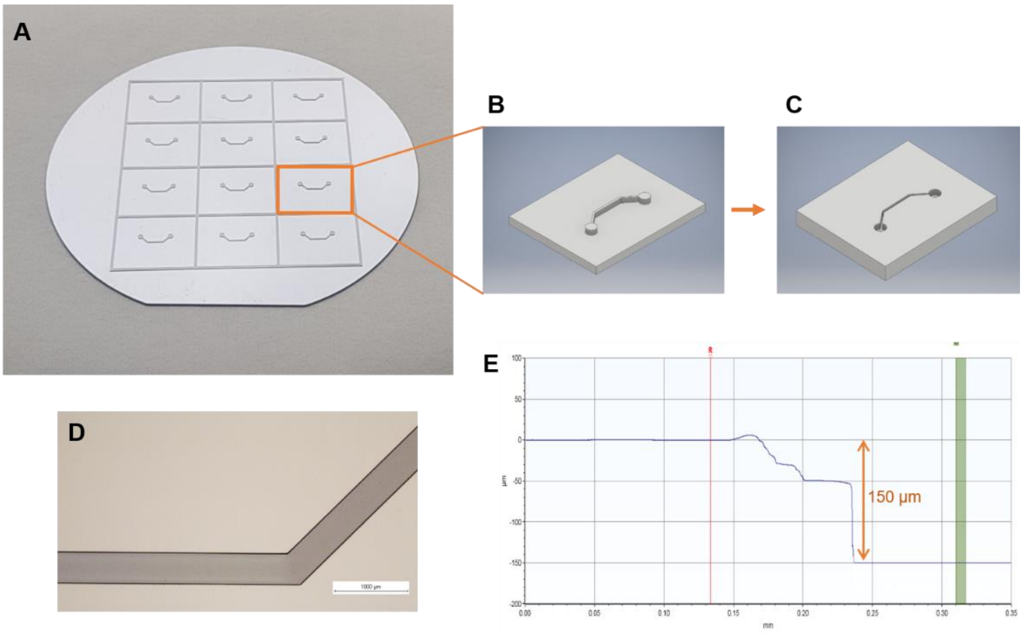


**Figure S1**. (A) Photographic representation of a SU-8 master mold fabricated by photolithography in this study. (B) Schematic representation of the patterns on the mold. (C) Schematic representation of the patterns on a PDMS layer, after using this mold for soft lithography. (D) Microscopy results to inspect the width of the channels. Scale bar: 1000 μm. (E) Profilometer results to inspect the height of the channels.

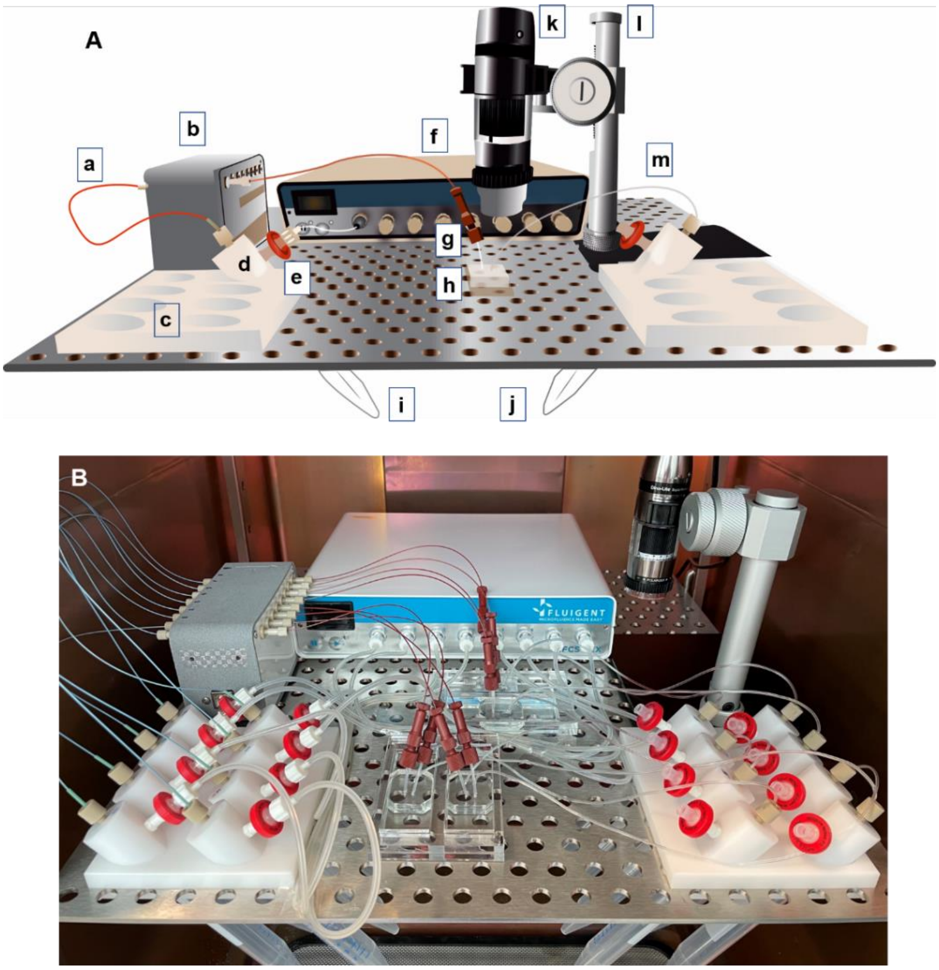


**Figure S2.** (A) Schematic representation of the microfluidic cell-culture setup used in this study, which consists of (a) tubing of the sensor (PEEK tubing), (b) flow sensor, (c) holder of the reservoirs, (d) caps for pressurization of the reservoirs, (e) filters to avoid contamination, (f) pressure pump, (g) micro-tight fitting to connect the sensor tubing and the chip tubing, (h) chip and chip holder, (i) media reservoir, (j) collection reservoir, (k) light microscope, (l) microscope stand, and (m) tubing of the chip (Tygon tubing). This schematic illustration depicts the connections, reservoirs, and tubing for only a channel of a chip. Each chip consists of two channels and the same connection method was applied to the other channel. The setup can be used to run eight channels (i.e. four chips), simultaneously. (B) Photographic representation of the microfluidic cell-culture setup used in this study. This setup was placed inside a cell-culture incubator and the pressure pump, flow sensor, and microscope were connected to a computer to control the system.
